# Simplifying the development of portable, scalable, and reproducible workflows

**DOI:** 10.1101/2021.04.30.442204

**Authors:** Stephen R. Piccolo, Zachary E. Ence, Elizabeth C. Anderson, Jeffrey T. Chang, Andrea H. Bild

## Abstract

Command-line software plays a critical role in biology research. However, processes for installing and executing software differ widely. The Common Workflow Language (CWL) is a community standard that addresses this problem. Using CWL, tool developers can formally describe a tool’s inputs, outputs, and other execution details in a manner that fosters use of shared computational methods and reproducibility of complex analyses. CWL documents can include instructions for executing tools inside software containers—isolated, operating-system environments. Accordingly, CWL tools are portable—they can be executed on diverse computers—including personal workstations, high-performance clusters, or the cloud. This portability enables easier adoption of bioinformatics pipelines. CWL supports workflows, which describe dependencies among tools and using outputs from one tool as inputs to others. To date, CWL has been used primarily for batch processing of large datasets, especially in genomics. But it can also be used for analytical steps of a study. This article explains key concepts about CWL and software containers and provides examples for using CWL in biology research. CWL documents are text-based, so they can be created manually, without computer programming. However, ensuring that these documents confirm to the CWL specification may prevent some users from adopting it. To address this gap, we created ToolJig, a Web application that enables researchers to create CWL documents interactively. ToolJig validates information provided by the user to ensure it is complete and valid. After creating a CWL tool or workflow, the user can create “input-object” files, which store values for a particular invocation of a tool or workflow. In addition, ToolJig provides examples of how to execute the tool or workflow via a workflow engine.

## Introduction

Software is fundamental to modern scientific research^1,2^. It can accelerate the pace of research, formalize algorithmic logic, and support reproducibility^3^. In a 2014 survey, 92% of academic scientists reported using software in their research^4^. Our article focuses on command-line software, which scientists use in many disciplines^5^ and which often provides advantages over point-and-click software, especially for computational reproducibility^3,5^. Command-line software includes tools that scientists have prepared for general use within the research community; but it also includes custom scripts that scientists create to analyze particular dataset(s). Popular languages for creating such scripts include Python, R, Perl, and bash. In this article, we describe processes for using command-line tools and scripts in research; we emphasize challenges associated with using them, combining them, and ensuring that others can use them; and we describe how *workflow engines* and *software containers* help to address these challenges. Our target audience includes bioinformaticians who wish to develop and deploy software tools. But we also demonstrate how workflow engines and software containers can be used to support reproducibility of research analyses, including those that use custom scripts.

A *workflow* is a defined series of steps for processing or analyzing research data^6^. Each step applies one or more computational tools to the data with specific inputs, outputs, and configuration settings. Defining these steps as a workflow provides advantages over using command-line tools alone. Most importantly, a workflow provides a vocabulary for describing how the tools are interconnected. It also provides more flexibility regarding the computing environments that can be used. For example, the Snakemake^7^ and Nextflow^8^ workflow engines facilitate execution of workflows on local-, cluster-, or cloud-based computers. This process is made easier when command-line tools are encapsulated in software containers^9^ because the tools are portable.

Different workflow-management systems use different methodologies for defining workflows and for interfacing with software used within those workflows^6^. This heterogeneity motivated creation of the Common Workflow Language (CWL), an open specification for describing command-line tools and workflows^10^. CWL documents are recognized by many workflow engines, including Snakemake^7^, *cwltool*, Toil^11^, Apache Airflow^12^, Tibanna^13^, and Arvados^14^.

It is possible to use workflow engines like Snakemake and Toil independently of CWL. It is also possible to use software containers on their own. However, by creating CWL documents, scientists can describe tools and workflows in a way that is standards-based and agnostic to the workflow engine(s) on which it is executed. CWL documents are merely text-based files and thus can be created manually via a text editor. Despite this simplicity, many researchers have yet to adopt CWL. Useful tutorials are available online^15^, but as CWL specification provides flexibility, researchers face a learning curve to ensure that documents are specified correctly. Some software applications are available to aid in this process. For example, *Rabix Composer* is a desktop application that enables researchers to create and edit CWL documents^16^. This application supports many features in the CWL specification and provides both text-based and visual editors. However, researchers unfamiliar with nuanced details of the CWL specification may find Rabix Composer too advanced for their needs, and it does not support the latest versions of CWL. Alternatively, CWL plugins exist for many code editors^15^. In addition, developers have created software packages that enable researchers to build CWL documents via application programming interfaces. However, many researchers who could benefit from CWL lack the programming expertise to use these resources.

In this article, we illustrate how to use CWL to describe command-line tools and workflows and to perform reproducible research analyses. We provide 27 examples of CWL documents for completing diverse types of research tasks, ranging from simple (e.g., printing custom messages to the console) to advanced (identifying differentially expressed genes or calling somatic mutations in cancer genomes). In addition, we introduce *ToolJig*, a Web application that enables researchers to create CWL documents.

*ToolJig* provides a simple, interactive interface that requires no installation and includes prompts to guide the user. Via *ToolJig*, a researcher can specify details about a tool’s expected inputs and outputs, operating-system environment, and auxiliary files (e.g., scripts, configuration files). Researchers can also create workflows that integrate these tools. *ToolJig* checks the information provided by the user to ensure it is complete and valid. After successfully describing a tool or workflow, the researcher can download CWL files and use *ToolJig* to create “input-object” files, which store input values for a particular invocation of a tool or workflow. In addition, *ToolJig* provides examples of how to execute the tool or workflow via a workflow engine.

### Using containers to manage software installation and configuration

First, we address software installation and configuration, which are essential steps to creating CWL tools and workflows. These seemingly simple steps are fraught with challenges. Although the software may be downloadable from a public website, installation instructions are sometimes vague, the process may involve many steps, and these steps may differ for each operating system. Such challenges led computational biologist Ian Holmes to quip, “You can download our code from the URL supplied. Good luck downloading the only postdoc who can get it to run”^17^. Package managers, such as *bioconda* and *bioconductor*, have helped to ameliorate these challenges, providing mechanisms to install software dependencies and track versions. These package managers function in a way that is mostly agnostic to the user’s operating system^18,19^. However, some software tools are not available via package managers, package managers depend on operating-system components that cannot be installed using the package managers themselves, and package managers may not guarantee that older versions of software and their dependencies remain available^3^. Software *containers* have gained popularity among scientists in recent years because they help to overcome these limitations.

Software containers provide a mechanism to encapsulate specific versions of software and their dependencies in a fully configured, operating-system environment^9^. Here we focus on the *Docker* ecosystem, which is commonly used for building, managing, and sharing software containers^9,20^. Other containerization systems are also available^21–25^; these systems are typically compatible with Docker. In academic computing environments, such as university-run cluster computers, Singularity has gained popularity because containers can be executed without administrative privileges.

Steps for configuring the operating-system environment and installing software within a container are documented in a “Dockerfile”. Using such a file, researchers can build a container *image*, a layered set of operating-system components. Commonly, the base layer is a minimal implementation of a Linux distribution (for example, Debian 10.3 or Ubuntu 18.04). Subsequent layers consist of software dependencies, configuration files, environment variables, etc. Once a container image has been created, it is portable and immutable. This is advantageous for computational reproducibility because one researcher can share an image with another researcher and know that its contents have remained static. A software *container* is an actively executing instance of a particular container image. Multiple containers of the same image can be executing simultaneously on the same computer (or different computers). Docker containers are always Linux-based; this is convenient for biology research because bioinformatics software is predominantly designed for Linux. Even though a container is Linux-based, it can be executed on non-Linux operating systems, such as Windows 10 or Mac OS 10, via a container *engine*. Container engines use virtual machines to facilitate this interaction^3^.

In addition to packaging scientific software, container images can package analysis code. For example, upon analyzing a given dataset, a researcher may wish to share the code with the research community. Many researchers post analysis code on websites such as GitHub (https://github.com) or Open Science Framework^26^. This practice can enable others to verify and reuse the code; it also benefits the original researcher who otherwise might lose track of analysis details^3^. But even when analysis code resides in the public domain, third-party researchers may experience difficulty executing it. Scripting languages like Python^27^ and R^28^ require interpreters. Analysis scripts may depend on specific versions of interpreters, but the third-party researcher may have a different version on their computer. In addition, most analyses rely on ancillary software packages. Such packages provide logic for parsing a certain type of file, performing statistical tests, creating graphics, etc. Versioning is critical: older (or newer) versions of software packages may be incompatible with the analysis code. Researchers can facilitate reproducibility by providing container images that include specific versions of any script interpreters or software packages that are necessary to execute an analysis.

*Binder* facilitates the containerization process for analysis code stored in GitHub repositories^29^. To use Binder, a researcher creates a configuration file that indicates which software is needed in the container image. For Python and R analyses, these configuration files indicate packages that must be installed, as well as version information. In other cases, a Dockerfile can be used to configure the environment more flexibly. After the researcher places the configuration file (and analysis code) in a GitHub repository, other researchers can visit the Binder website and re-execute the analysis. Behind the scenes, Binder provisions a cloud-based computer and executes the code within a container. This solution is effective for relatively short-running analyses that require modest computational resources, that use small datasets, and that are ready to be released publicly. However, many analyses do not meet these criteria.

Longer, more data-intensive analyses can be executed via workflow-management systems.

A *workflow* is a defined series of steps for processing or analyzing research data^6^. Each step applies one or more computational tools to the data with specific inputs, outputs, and configuration settings. Defining these steps as a workflow provides advantages over using command-line scripts alone. Most importantly, a workflow provides a vocabulary for describing how the tools are executed and interconnected. It also provides more flexibility regarding the computing environments that can be used. For example, the Snakemake^7^ and Nextflow^8^ workflow engines facilitate execution of workflows on local-, cluster-, or cloud-based computers. This process is made easier when command-line tools are encapsulated in container images because the tools are portable.

### Basic elements of a Common Workflow Language tool description

To describe execution of a command-line tool, a researcher creates a text-based file according to the CWL Command Line Tool Description specification^30^. CWL files can be structured using either the YAML or JSON data-serialization formats^31,32^. Here we provide an overview of key components of CWL tool descriptions.

A CWL tool describes inputs that will be used when the command-line tool is executed. A data type (or schema) should be defined for each input, indicating whether it represents a string, number, Boolean value, file, directory, or array (a data structure with multiple values). In practice, these inputs are generally data files and configuration settings for the software. These definitions help users of the tool understand the nature of each input and make it easier for inputs to be validated. For example, if a command-line tool expects a particular input value to be an integer (e.g., number of threads), a workflow engine can verify that the user has specified an integer before executing the tool.

After inputs have been defined, a tool description must provide instructions for constructing a command from the inputs. These commands can be based on a sorted ordering of the inputs. Alternatively, the researcher can specify a template for the command, using placeholders for the inputs. Such templates can represent either a single command or multiple commands.

As a tool executes, it can generate three types of outputs that might be useful to a researcher: 1) standard output, 2) standard error, and 3) new files. *Standard output* often consists of informational messages printed to the console; but it may also contain data to be used as input for another tool. *Standard error* typically consists of errors, warnings, or diagnostic information printed to the console. Many command-line tools produce new data files that have resulted from execution of the tool. A CWL tool description must indicate which of these outputs are expected so that a workflow engine can “collect” them after executing the tool.

### CWL tool examples for printing simple command-line messages

In the GitHub repository that accompanies this article, we have provided example tool descriptions, formatted according to the CWL specification (https://github.com/srp33/ToolJig/tree/master/examples). The first series of examples is stored in the `hello` subdirectory.

The first example, `01_hello.cwl` requires two inputs: 1) a person’s given name and 2) the person’s surname. It uses the `baseCommand` field to construct a command based on ordinal positions specified for the inputs; the resulting command prints a greeting for that person. In this simple example, the only output is the greeting sent to standard output, which is redirected to a file called `01_output.txt`. In the GitHub repository, `hello_objects.yml` is an example input-object file for this tool. It species a person’s given name (“Fernanda”) and surname (“Dantas”). The user could execute the tool via the *cwl* workflow engine using this command: `cwltool 01_hello.cwl hello_objects.yml`. The output would be, “Hello, Fernanda Dantas”.

Suppose we wished to alter the greeting to include an exclamation point and to indicate the person’s age. First, we would add an input for the person’s age (an integer). Second, we would need to update the command that will be executed. However, CWL’s `baseCommand` field provides limited flexibility for constructing commands. Instead, our example provides a command template using the `ShellCommandRequirement` and `arguments` fields and uses placeholders within the template for each input. As shown in `02_hello.cwl` (and Figure 1A), we use the following template: `echo Hello, $(inputs.given_name) $(inputs.surname)! You are` `$(inputs.age) years old.`

**Figure 1:**
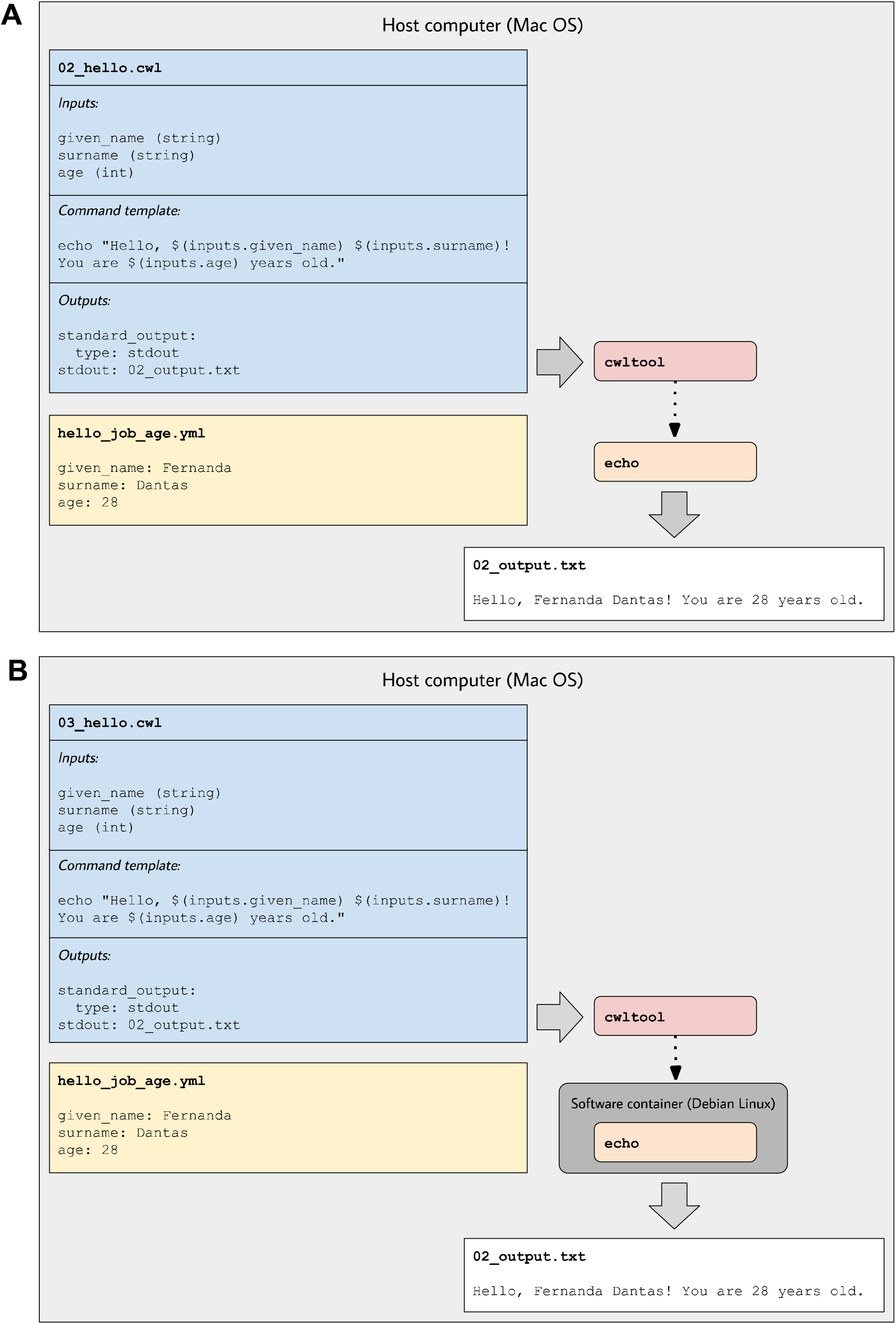
Illustration of tool descriptions for printing simple greetings. In the examples associated with this article, we provide tool descriptions that illustrate how to print custom greetings at the command line. These diagrams illustrate the `02_hello.cwl` (**A**) and `03_hello.cwl` (**B**) examples. In **A**, the tool description indicates which inputs that must be specified, along with a template for executing the command; it also indicates that a message will be printed to standard output and that this message should be stored in a file called `02_output.txt`. The `hello_objects_age.yml` input-objects file stores values for a particular invocation of the tool. In **A**, the `cwltool` workflow engine uses the host computer’s operating system to execute the tool; thus the echo command must be supported on that operating system. In **B**, the tool description defines a software container environment; thus `cwltool` executes the command within a container, which provides the echo command (packaged with the Debian Linux operating system).

We use a `$` character and parentheses to indicate placeholders for input variables. We prefix each input variable with “inputs.” to indicate that they will be specified as inputs.

By default, the first two examples would be executed within the same operating-system environment as the workflow engine. Accordingly, these tools can only be executed on operating systems that support the `echo` command. Many commands are only supported on particular operating systems—or their behaviors differ by operating system. So in `03_hello.cwl` (and Figure 1B), we use a Docker image based on the “buster” release (version 10.3) of the Debian Linux operating system. The `DockerRequirement` field is added (in this case, two lines of text). Before executing the tool, a researcher would install a container engine such as Docker Desktop. Then, when executing the tool, the workflow engine would integrate itself with the container engine, which would identify any input files or directories and create container *volumes* so that the files or directories could be accessed from within the container. With this option enabled, the command-line tool becomes portable—it can be executed on any computer that supports the container and workflow engines.

### CWL tool examples for performing simple data analyses

The second series of examples is stored in the `examples/bmi` subdirectory on the GitHub site. The `01_bmi.cwl` tool provides a simple example of a reproducible, quantitative analysis that could be performed using CWL. It accepts as input a tab-separated file containing names, weights, and heights of (fictional) individuals. A second input specifies the name of the column in the tab-separated file that contains weight information. The third input specifies the column name containing height information. The fourth input is the name of an output file that will be created. This example illustrates the use of an

*auxiliary file*. Under the `InitialWorkDirRequirement` field, the contents of a Python script are stored. This script is used to calculate Body Mass Index (BMI) values for each person in the input file and store those values in a *BMI* column in the output file. Below is the command template that we use.

~~~
python calculate_bmi.py “$(inputs.input_file.path)”
“$(inputs.weight_column_name)” “$(inputs.height_column_name)”
“$(inputs.output_file_name)”
~~~

The command template specifies the inputs as arguments to the Python script. When a file input is used, the workflow engine stores metadata about the file in an object with multiple attributes. Thus to reference the file’s path within the command template, we append “.path” to the input name. As the workflow engine executes the tool, it stores the auxiliary Python script inside the container, invokes the script, and collects the output file that the script generates.

If a researcher wished to ensure that others could reproduce the BMI calculations, they would only need to share the CWL file, the input-object file (`01_bmi_objects.yml`), and the data file (`biometric_data.tsv`). However, many analyses use data stored in online repositories. In such cases, it is convenient for a CWL tool to pull data directly from an online repository. The `02_bmi.cwl` tool description and Figure 2 illustrate this approach. Similar to the previous example, it extracts names, weights, and heights from a tab-separated file and adds a BMI column. However, it pulls the file from a web server (in this case, our GitHub repository). The command template is similar to the previous example. Again, we use a software container based on Debian Linux. But this time, we use the `NetworkAccess` field to enable the container to connect to external computers. The tool emits messages to both standard output and standard error; these messages are stored in files called “02_output.txt” and “02_error.txt”, respectively.

**Figure 2:**
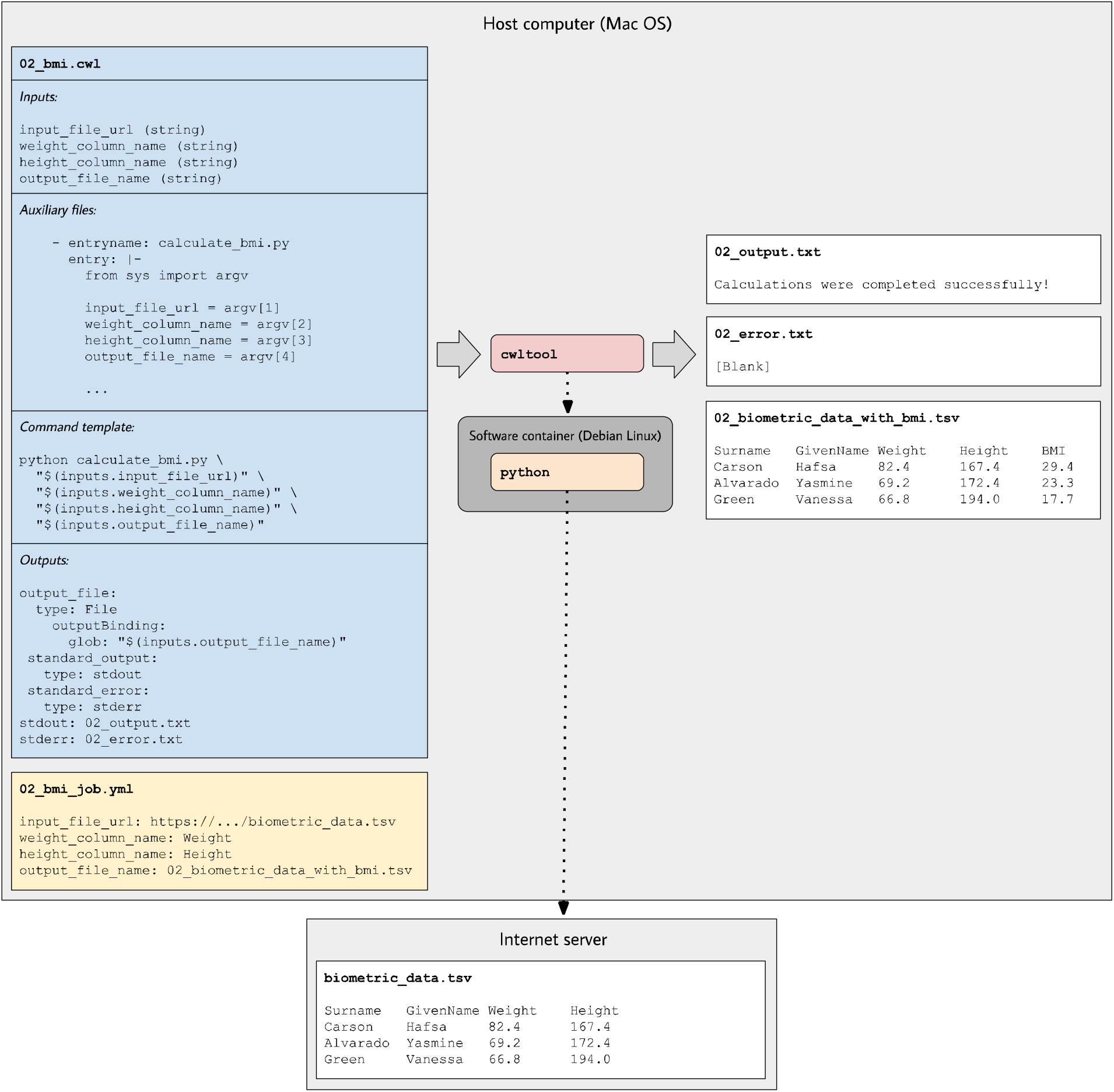
Illustration of tool descriptions for calculating individuals’ body mass index. In the examples associated with this article, we provide tool descriptions that illustrate how to calculate body mass index (BMI) values based on individuals’ weights and heights stored in a tab-separated-value file. This diagram illustrates the `02_bmi.cwl` example. The tool description indicates the expected inputs. In this case, the URL of a data file must be provided. That file must contain a column that stores weights (in kilograms) and a column that stores heights (in centimeters). In the input-objects file (`02_bmi_objects.yml`), the user specifies the names of these columns. The final input is the name of an output file that will be generated. This file will store the original data and a new column with the calculated BMI value for each individual. As the tool executes, python (within a software container) downloads the input file, performs the calculations, generates the output file, and stores standard output and standard error in text files.

### CWL tool examples for processing transcriptomic data

The third series of examples (`examples/transcriptomic` subdirectory) are wrappers around existing tools for processing transcriptomic data. In both cases, we use R packages from the Bioconductor suite^19^. Although R and Bioconductor are designed to be compatible with all major operating-systems, some packages require dependencies that are operating-system specific. Furthermore, many Bioconductor packages provide a large number of functions and options. Researchers can create CWL tool descriptions that install dependencies (within a container image) and support a narrower range of options. The researchers might then share this tool with other researchers, enabling them to apply the tool more easily to their own data. Alternatively, they might use the tool as a way to support reproducibility for their own analyses.

The Single-Channel Array Normalization (SCAN) algorithm normalizes data from gene-expression microarrays, correcting for background noise and oligonucleotide-binding biases^33^. The SCAN method is implemented in the SCAN.UPC package in Bioconductor. It can download data directly from Gene Expression Omnibus (GEO)^34^. The `scan_normalize.cwl` example illustrates how this functionality could be incorporated into a CWL tool. The base container image in this example includes the core Bioconductor components; our Dockerfile extends this image by installing the SCAN.UPC package. In addition, our example uses an auxiliary file containing R code that invokes the SCAN function within this package to normalize a given GEO series. Upon executing, this tool produces a tab-separated output file containing normalized measurements for all biological samples in the series. The tool could be customized further, for example, to perform gene-level rather than probeset-level summarization or to perform a quality-control analysis.

Commonly, researchers seek to identify genes that are differentially expressed between two conditions. The DESeq2 package is popular for performing such analyses with RNA-Sequencing data^35^. Our `deseq2.cwl` example illustrates how this process could be incorporated into a CWL tool. Similar to the previous example, this tool uses a container image with Bioconductor core components and installs the “DESeq2” package. In addition, it installs the readr and dplyr packages^36,37^, which we use to read and parse the data before identifying differentially expressed genes. The first two inputs are URLs to data files containing gene counts and phenotype information in tab-separated formats. The third input is a string representing a design formula; users of the tool can customize the differential-expression calculations based on the dependent variable of interest as well as any covariates. In the example input-object file (`deseq2_objects.yml`), we use data from an RNA-Sequencing experiment that compared two inbred mouse strains commonly used for neuroscience research^38^; the data had been aligned to a reference genome, and gene counts had been quantified previously.

### CWL workflow examples for performing mathematical calculations

The examples so far have illustrated how to execute command-line tools in isolation. CWL workflows must specify at least one input(s) and one output(s) for the entire workflow. In addition, the researcher must define steps that each consist of a tool with input(s) and output(s). The researcher indicates whether each step’s input should be populated by an input for the entire workflow or by the output of a previously completed step. The workflow’s output(s) consist of the output(s) of one or more of the steps. As with CWL tools, the researcher must create an input-object file that provides input values for a particular execution of the workflow. Upon executing the workflow, the workflow engine evaluates the sequence of steps that must be executed and connects inputs with outputs, as needed.

We provide three example workflows in the `examples/workflows/math` subdirectory of the GitHub repository. The `add_sqrt_workflow.cwl` example accepts two integers as inputs, sums them, calculates the square root of the sum, and then stores the square root of the sum in an output file. This example illustrates the basic process of using an output from one tool as input to another. The `recursive_sqrt_workflow.cwl` example reads a number from a file, calculates the square root of that number, calculates the square root of the resulting number, and saves the output to a file. This workflow demonstrates the ability to invoke the same tool recursively. The `secondary_sqrt_workflow.cwl` example calculates the square root of a number stored in a file and saves the result to an output file. It does the same for two secondary files. It then sums those values and writes the sum to a file. This example demonstrates using secondary files within a workflow. Secondary files are commonly used in genomics and simplify the process of working with groups of files that are necessary to complete a particular task.

### CWL workflow example for identifying somatic variants in a cancer genome

The examples in the `examples/workflows/somatic` subdirectory demonstrate a process for call somatic variants from Illumina sequencing reads. We use paired-end reads from tumor and normal cells for an individual from the Texas Cancer Research Biobank^39^. (Although these data are publicly available, they are subject to some data-use restrictions^39^.) To shorten execution times, we use a subset of the data: the first 10,000,000 reads from lane 2 of the sequencing run. Furthermore, our analysis is limited to essential steps for preparing the data and calling variants. Additional steps like annotation and filtering would improve sensitivity and specificity of the variant calls; accordingly, researchers should interpret our variant calls with caution.

### In these examples, somatic-variant calling occurs in a series of steps (Figure 3)

1. *Download and index a human reference genome (version hg38)*. We use the Linux *wget* and *gunzip* utilities to download and decompress a FASTA file from the UCSC genome repository^40^. We also use bwa, samtools, and Picard Tools to index the FASTA file and create a sequence dictionary^41–43^.
2. *Preprocess reference files containing known polymorphic sites in preparation for base-quality score recalibration (BQSR)*. We download Variant Call Format files^44^ from the Genome Analysis Toolkit (GATK) resource bundle^45^ and use a custom Python script to adjust chromosome identifiers that may differ across reference genomes. We also use Picard tools to sort the reference files.
3. Download the raw sequencing reads (FASTQ files). The files are stored in a publicly available, Open Science Framework repository^26^.
4. Trim adapter sequences and low-quality bases using atropos^46^.
5. Align the trimmed reads to the reference genome using *bwa mem*^41^. A read-group string is also specified.
6. Sort and index the resulting BAM files^42^ using sambamba^47^.
7. Mark duplicate reads and re-index the resulting BAM files using sambamba.
8. Derive a BQSR table from each BAM file using GATK.
9. Apply the BQSR table to the BAM files using GATK.
10. Call somatic single-nucleotide variants and small insertions/deletions using Mutect2^48^. This produces a VCF file.
11. Call somatic structural variants using Delly^49^. This produces a VCF file.

This workflow executes many of the same steps for a normal DNA sample and a tumor DNA sample. These steps are independent of each other. Accordingly, when multiple computing cores are available, the workflow engine may execute these steps in parallel.

Most of the tools in this workflow use container images from the BioContainers project, which provides thousands of Docker images that encapsulate biology-related software^50^. In some cases, we used a BioContainers image that had been built for a specific version of a software package in the Bioconda project.^18^ In cases where we used multiple packages for a given task, we started with a base image from Biocontainers and used Bioconda to install the packages. In the case of GATK, we used container images that had been created by the Broad Institute and stored on Docker Hub.

In most cases, we followed the recommendation that a single container image use only a single software package^51^. However, in some cases, we determined that it was more sensible to use multiple software packages in a given CWL tool. For example, when preparing index files for the reference genome, we use three separate software packages. In contrast, sometimes we used the same software in multiple CWL tools. For example, the BQSR steps are computationally intensive; thus, we separated them into distinct CWL tools so that computational resources can be allocated at a more granular level. In this sense, each CWL tool represents a practical unit for data processing, not necessarily a particular software package.

## ToolJig

### Using ToolJig to create CWL tool descriptions

In manufacturing, a “jig” is used by toolmakers to ensure that products are created in a repeatable, consistent pattern. Similarly, ToolJig provides a means to generate CWL tool descriptions, workflows, and input-object files in a repeatable, consistent manner. ToolJig is a Web application that uses the Vue.js framework^52^. Its functionality is divided into two pages: one for creating CWL tools and one for creating workflows. It is available from https://srp33.github.io/ToolJig/. To create a tool description in ToolJig, users specify the following:

1. A unique identifier. This identifier is used in the name of the CWL file that is generated, as well as for tagging the Docker image.
2. A short label that describes the tool’s purpose and function.
3. Optionally, a longer description that provides more detailed documentation about the tool.
4. Dockerfile contents. These instructions indicate the base container image that should be used and any additional commands necessary to build and configure a container image for the tool. The tutorial by Nüst, et al. provides helpful recommendations on authoring Dockerfiles^53^.
5. Author information. Optionally, the tool author can specify their name and ORCID identifier^54^. This information helps to ensure that authors are credited for their work.
6. Software license. The tool author can select from among 7 popular licenses, thus indicating conditions under which others can use the CWL document. This license may or may not be identical to the license specified for the software itself.
7. Inputs. Users specify a name, type, and description for each input. Supported types are integer, string, File, and “Output File.” The File type allows the user to indicate that an input file is expected and asks the user to specify the EDAM format of the file^55^. Additionally, input files may be associated with secondary files. For instance, as our examples illustrate for somatic variant-calling, BAM files must be accompanied by index files. Rather than specify these as two separate inputs, we indicate that an index file is secondary to a BAM file. “Output File” is a convenience type that is used when a tool author wants users to be able to specify the *name* of an output file that will be generated as the tool executes. Because this requires user input, ToolJig provides it as an input option. When a tool author specifies this type, ToolJig creates a *string* input for the file name, along with a corresponding output element, thus simplifying this process for the user.
8. Auxiliary files. The tool author can enter the name and contents of any auxiliary files that will be used. These tools are commonly programming scripts and can be written in any programming language that the Docker image supports.
9. Command template. The tool author specifies a template for executing the tool at the command line. Each input must be specified at least once in this template; ToolJig provides syntax suggestions to the user. ToolJig uses these command templates as an alternative to the `baseCommand` field. As our examples illustrate (Figure 4), command templates provide flexibility in the way that commands are constructed, including the use of multiple commands. They provide the additional benefit that inputs do not have an inherent order and thus can be specified in the command template in any order (and multiple times, if desired).
10. Outputs. Aside from any “Output Files” that may have been specified as inputs, the user may declare additional output files. For instance, the `somatic/trim_fastq.cwl` example specifies that trimmed FASTQ files should have the same names as the corresponding input

**Figure 3:**
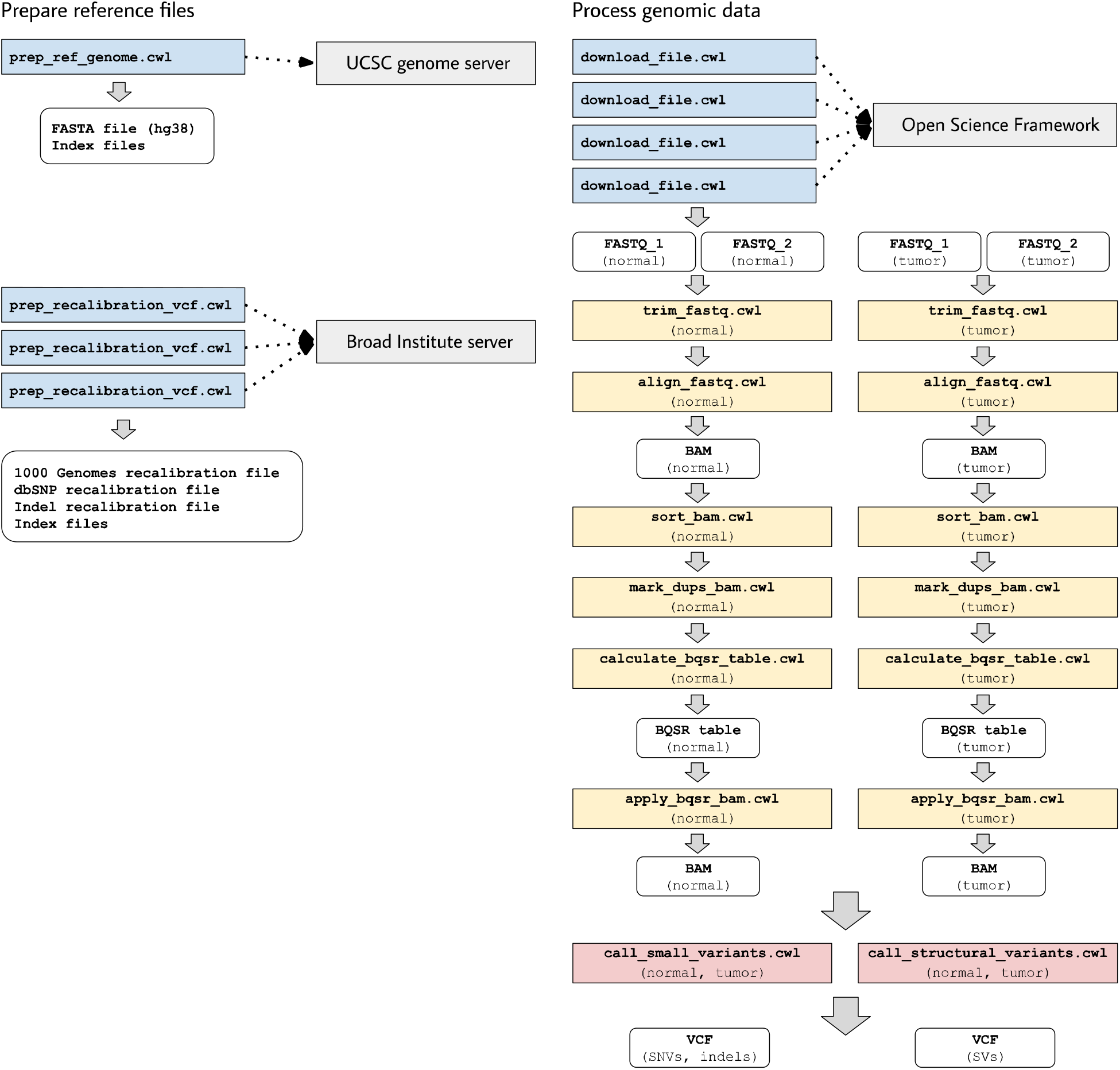
Illustration of tool descriptions for calling somatic variants from a cancer genome. In the examples associated with this article, we provide tool descriptions that illustrate how to call somatic variants from second-generation sequencing data for a cancer genome (compared against a normal genome from the same patient). This process requires execution of 11 distinct tools in a defined succession of steps (a workflow). Two tools (prep_ref_genome.cwl and prep_recalibration_vcf.cwl) prepare reference files associated with a given human reference genome. These tools download data files from public Internet servers and then create index files and standardize contig identifiers. The third tool (download_file.cwl) downloads FASTQ files from an Internet server. The remaining tools process the normal and tumor sequences separately before comparing the tumor genome against the normal genome to identify single nucleotide variants, indels, and structural variants.

**Figure 4:**
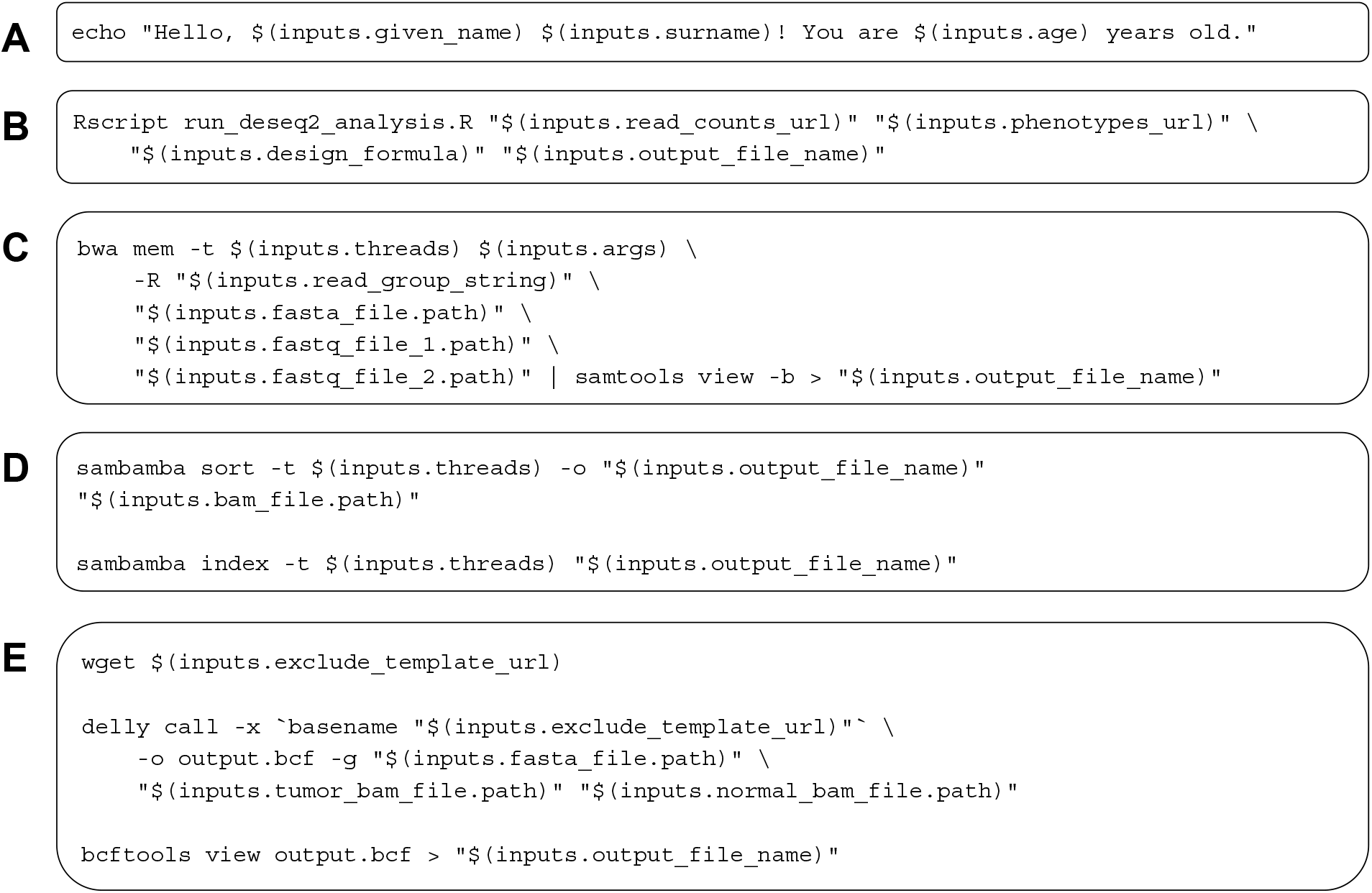
Examples of command templates used in CWL tool descriptions. These examples illustrate diverse types of command templates for configuring execution of CWL tools. In each example, placeholders are used for inputs. When the tools are executed, the placeholders are replaced with input-objects values. **A)** A simple greeting is printed to standard output. **B)** An R script (stored as an auxiliary file within the tool description) is executed; this script performs a differential-expression analysis using the DESeq2 package. **C)** The *bwa* software aligns FASTQ files to a reference genome and pipes the output to *samtools*; the output is then converted to BAM format. This example illustrates a scenario in which two complementary software packages are used to perform a data-analysis task. Although these packages could be incorporated into distinct CWL tools, we combine them because read alignment and BAM conversion are typically performed jointly. **D)** The *sambamba* software sorts and then indexes a BAM file. **E)** The *Delly* software identifies structural variants in a cancer genome. *Delly* can be configured to exclude telomere and centromere regions as well as unplaced contigs. This example downloads an exclusion file, invokes *Delly*, and converts the output to VCF format. Examples **D** and **E** illustrate additional scenarios in which related tasks are executed as practical units.

#### FASTQ files. To indicate this, the user specifies a CWL-based expression

`$(inputs.fastq_file.basename)`. Additionally, if the user wishes to collect standard output or standard error, they may specify the names of files that will store these messages.

After a user has specified all required elements, ToolJig generates a YAML document that conforms to the CWL specification; the user may download this document. ToolJig also generates a form in which the user can indicate a value for each input. Subsequently, the user can download an input-object file in YAML format.

### Using ToolJig to create CWL workflows

When using ToolJig to create a workflow, researchers first enter metadata: a unique identifier, label, description, author information, and software license. Subsequently, the researcher uploads at least one tool description. The researcher then defines two or more workflow steps. For each step, they specify a unique name and the tool that will be used in that step. For each of the tool’s inputs, the researcher indicates whether the input will be populated from the output of a preceding step or from a workflow input. The user may also indicate that any of the tool’s outputs will become outputs for the overall workflow. As with tool descriptions, ToolJig validates the user’s input and then generates a CWL document and input-object file that can be downloaded.

## Discussion

Progress in biology research is hindered when software tools are difficult to install, when inputs and outputs are inadequately or inconsistently specified, and when it is difficult to combine tools into workflows. The CWL specifications—in combination with package managers and software containers— are helping to alleviate these longstanding challenges. Moreover, CWL tool and workflow descriptions can facilitate reproducible research. Rather than simply providing analysis code alongside journal manuscripts, researchers can provide CWL documents. As illustrated in our examples, CWL documents provide instructions for executing analyses in software containers that encapsulate all relevant dependencies (Figure 5), along with ancillary scripts and instructions for accessing data files. People who read (or review) the manuscripts can then repeat the analyses, without needing to install any software other than a relevant workflow engine and container engine, even if their operating system or configuration differs from the authors’.

**Figure 5:**
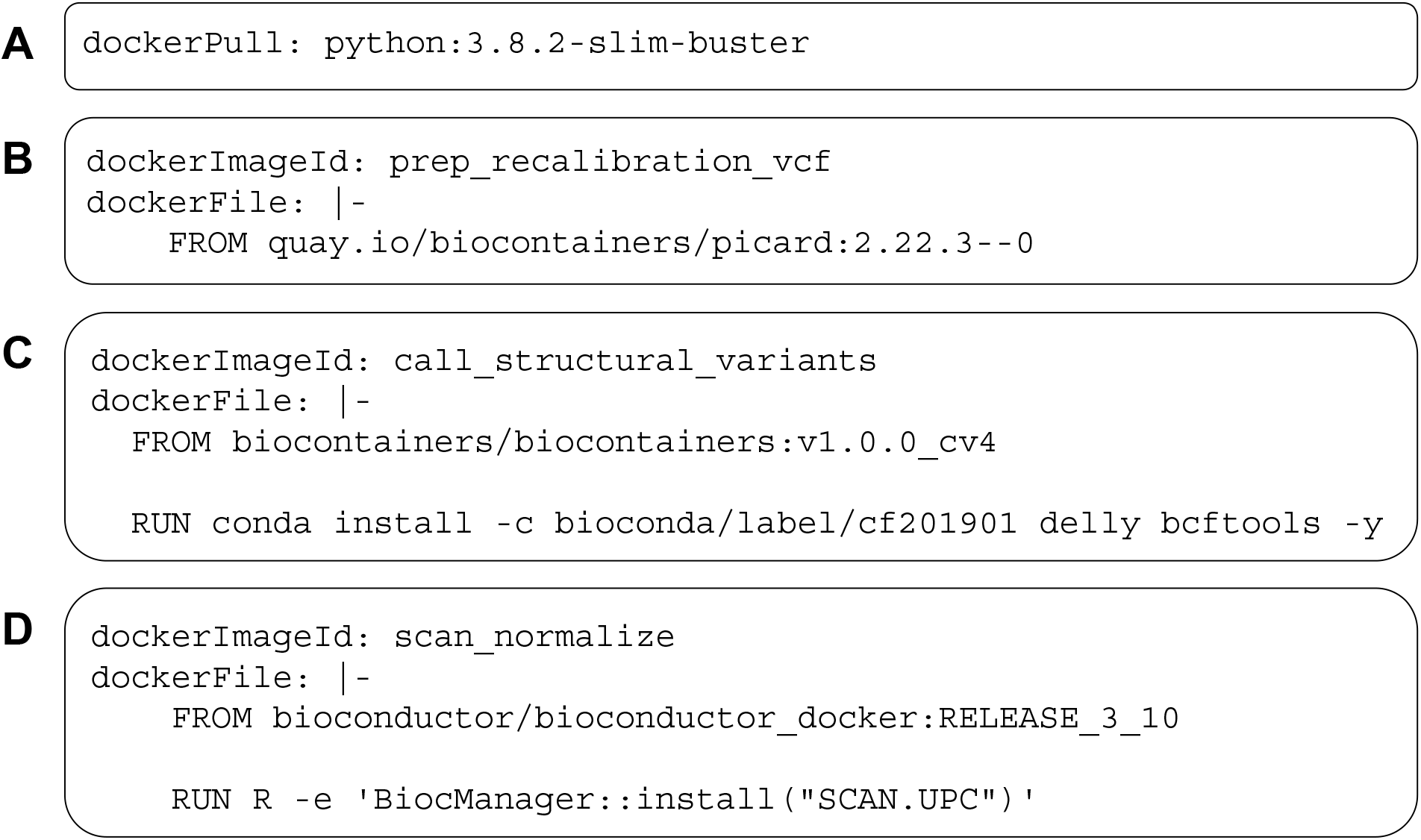
Examples of DockerRequirement specifications used in CWL tool descriptions. These examples illustrate diverse ways to configure CWL tools to be executed in software containers. In **A**, a container image is pulled from Docker Hub; this image encapsulates a minimal (“slim”) version of Debian Linux 10.3 (“buster”) and includes the Python 3.9 interpreter. In **B**, the contents of a Dockerfile are included within the CWL description. In this case, the Dockerfile is simple—it pulls an existing image from quay.io. This image is provided as part of the Biocontainers project and includes the Picard Tools software. **C** uses a base image from Biocontainers and the Bioconda package manager to install the *Delly* and *bcftools* software within the image. **D** uses a base image from Bioconductor and executes R code to install the SCAN.UPC normalization package within the image.

Multiple online repositories provide CWL documents, including the Dockstore tool registry^56^ and GitHub. A search on GitHub for CWL documents that use the FastQC software^57^, for example, resulted in 616 matches (May 15, 2020). Researchers can reuse and adapt these documents. However, in cases where reuse is infeasible or extensive adaptations are necessary, scientific progress may be accelerated as researchers, including non-bioinformaticians, gain greater efficiency in creating CWL documents. ToolJig aims to facilitate this process, enabling researchers to build CWL tools interactively, without needing to gain a deep understanding of the CWL specification or YAML syntax.

One advantage of CWL is that it can be used with diverse workflow engines. Whether or not they support CWL, most workflow engines provide custom languages or programming interfaces for creating workflows. Relatively little support is available for migrating from these engine-specific solutions to CWL in an automated manner. However, when these engines support execution within Docker-compatible containers, researchers can migrate these tools manually using ToolJig (or other means).

Providing better support in existing workflow engines for exporting to CWL will be a positive step toward ensuring that CWL truly becomes a common language for command-line tools and workflows.

The CWL specification provides considerable flexibility for describing command-line tools and workflows. Our goal was to support common use cases for biology research. For the sake of simplicity and to reduce barriers of entry for new creators of CWL documents, we deliberately do *not* support some optional features within the CWL specification. These include input directories, dependent and exclusive parameters, process requirements, hints, and output directories. To learn about augmenting CWL documents with these features, researchers can consult the CWL specifications.

ToolJig has no dependencies other than a modern Web browser. Accordingly, it can be used from virtually any computer with no installation process. When ToolJig is updated, the user simply needs to refresh their browser. A tradeoff to this simplicity is that ToolJig does not provide a direct means of testing tools or workflows. However, the `cwltest` utility provides a command-line testing framework, enabling researchers to compare tool and workflow outputs with expected results. Researchers implementing CWL in production systems would benefit from using such a utility for validation.

## Conclusions

CWL documents can formalize execution of command-line tools and workflows. We have summarized the key components of CWL tool descriptions and provided examples to illustrate these concepts. In addition, we have described ToolJig, a Web application that enables researchers to create CWL documents interactively. We hope these resources will benefit researchers from diverse backgrounds to more easily create CWL documents and thus advance sharing of methods and computational reproducibility.

## Declarations

### Ethics approval and consent to participate

Not applicable. *Consent for publication* Not applicable.

### Availability of data and material

The source code and examples are located at https://github.com/srp33/ToolJig.

### Authors’ contributions

SRP, ECA, and ZEE conceptualized the ToolJig application and generated prototypes. SRP and ECA created the application. SRP and ZEE prepared the tool examples. JTC and AHB provided critical feedback on the application’s functionality. SRP wrote the manuscript. All authors provided critical feedback on the manuscript.

We acknowledge the Texas Cancer Research Biobank and Baylor College of Medicine Human Genome Sequencing Center for providing openly available cancer-genome data used in our examples.

